# Establishing betaxanthin pigment biosynthesis in cyanobacteria

**DOI:** 10.1101/2025.06.16.659934

**Authors:** Sayali S. Hanamghar, David A. Russo, Silas Busck Mellor, Julie A. Z. Zedler

## Abstract

Betalains are water-soluble pigments with two major classes: red-violet betacyanins and yellow-orange betaxanthins. These pigments are increasingly being sought after as natural replacements for synthetic pigments in the food industry. Traditionally, betalains are extracted from cultivated plants but, due to low native pigments, the process is inherently inefficient. Now, as consumer demand increases, scalable and sustainable production routes are needed. To address this challenge, we engineered a heterologous pathway for the production of betaxanthins into cyanobacteria. The pathway consists of the cytochrome P450 CYP76AD1 and the L-DOPA 4,5-dioxygenase DODA1 from *Beta vulgaris* (beet). Introduction of the two-enzyme betaxanthin pathway in *Synechocystis* sp. PCC 6803 did not result in detectable betaxanthins. Subsequent metabolic adjustments to the shikimate pathway, using a feedback resistant AroG^fbr^ from *E. coli*, led to an overaccumulation of the aromatic amino acids phenylalanine, tryptophan, and tyrosine, and the production of 0.11 mg L^-1^ phenylalanine-betaxanthin. Optimization of the cultivation conditions (i.e., growth in nutrient-rich medium and CO_2_-enriched air) increased titers almost 150 times and led to the production of phenylalanine-betaxanthin with a final titer of 15.3 mg L^-1^. Our work establishes a microbial system for photoautotrophic betaxanthin pigment production without the need for exogenous amino acid supplementation.

## 1. Introduction

Betalains are nitrogen containing, water-soluble, red-violet and yellow-orange pigments. They are found mostly in plants of order Caryophyllales such as *Beta vulgaris* (Timoneda et al., 2019), *Amaranthus tricolor* (Chang et al., 2021) and *Dianthus caryophyllus* (Sadowska-Bartosz and Bartosz, 2021). Betalains have also been found in funghi (Babos et al., 2011; Hinz et al., 1997)and the bacterium *Gluconacetobacter diazotrophicus* (Contreras-Llano et al., 2019). Betalains have applications in the pharmaceutical and nutraceutical industries due to their antioxidant and anti-inflammatory properties. In addition, they are widely used in the food industry as a red food colorant (Sadowska-Bartosz and Bartosz, 2021).

Betalain biosynthesis starts with tyrosine. In the first step, tyrosine is hydroxylated to L-3,4-dihydroxyphenylalanine (hereafter L-DOPA) by a cytochrome P450, CYP76AD1 (Hatlestad et al., 2012). In the second step, 4,5-DOPA-extradiol-dioxygenase (DODA1) catalyses the ring-opening oxidation of L-DOPA to form an intermediate 4,5-seco-dopa that spontaneously cyclizes into betalamic acid, the common chromophore of betalains (Fernando Gandía-Herrero and García-Carmona, 2012). Betalamic acid can then spontaneously condense with amines and their derivatives to form betaxanthins. Betalamic acid can also spontaneously condense with cyclo-DOPA and/or its glucosyl derivatives to form betacyanins. Betaxanthins are yellow-colored pigments with absorption maxima between 460 and 480 nm that exhibit strong fluorescence due to the structural properties of the betalamic acid moiety when connected to an amine group (Spórna-Kucab et al., 2023). Betacyanins, on the other hand, are weakly fluorescent red and violet pigments with two absorption maxima; one in the UV range between 270 and 280 nm and other in the visible range between 535 and 540 nm (Azeredo, 2009).

Heterologous betalain biosynthesis has previously been established in crop plants (Grützner et al., 2021; Polturak et al., 2017; Wang et al., 2024) and heterotrophic microbial hosts such as *Escherichia coli* (Guerrero-Rubio et al., 2019; Hou et al., 2020) and *Saccharomyces cerevisiae* (Babaei et al., 2023; DeLoache et al., 2015). However, plant systems suffer from inherently low areal productivities and the existing microbial systems typically require feeding with amino acids or glucose to increase betalain titers. The aim of this study was, therefore, to establish the production of betaxanthins into a photoautotrophic microbial host. Cyanobacteria are prokaryotes that perform oxygenic photosynthesis. In recent years, there has been increasing evidence of the potential of cyanobacteria as whole cell catalysts (Jodlbauer et al., 2021; Toepel et al., 2023). This is particularly the case for industrially-relevant redox reactions that can take advantage of the excess reducing power generated by photosynthesis for light-driven catalysis (Russo et al., 2019a; Santos-Merino et al., 2021; Spasic et al., 2022; Torrado et al., 2022).

In this study we use *Synechocystis* sp. PCC 6803 (Syn6803) as a host to express *CYP76AD1* and *BvDODA1*, the two genes that constitute the biosynthetic pathway for betaxanthin production in *Beta vulgaris*. We show that sole expression of the betaxanthin pathway was not sufficient to produce betaxanthins to a detectable level. Addition of a feedback-resistant *aroG* (*aroG^fbr^*) increased the phenyalanine (Phe), tryptophan (Trp) and tyrosine (Tyr) pools and led to the detection of 0.1 mg L^-1^ of Phe-betaxanthin (Phe-Bx). Subsequent high-density cultivation improved Phe-Bx production almost 150 times to a final yield of 15.3 mg L^-1^. Our work provides the first proof of concept for the production of betalains in a photoautotrophic microbe.

## 2. Materials and Methods

### 2.1. Cyanobacterial strains and cultivation conditions

For this work a glucose-tolerant, non-motile, *Synechocystis* sp. PCC 6803 sub-strain ‘‘Kaplan’’ and *Synechococcus elongatus* PCC 7942 (obtained from the Pasteur Culture Collection of Cyanobacteria) were used. Cultures were maintained at 30 °C with approximately 50 μmol photons m^−2^ s^−1^ continuous illumination on 1.5% agar plates of BG-11 medium (Stanier et al., 1971a) supplemented with 5 mM 4-(2-Hydroxyethyl)piperazine-1-ethanesulfonic acid (HEPES) (pH 7.5) and 3 g L^-1^ sodium thiosulfate pentahydrate. Liquid cultures were grown in BG-11 supplemented with HEPES (BG-11_H_) in 50 mL tissue flasks (Greiner, product no.: 690175) at 30 °C with continuous illumination (approximately 50 μmol photons m^−2^ s^−1)^ and shaking at 125 rpm. Engineered strains were supplemented with 50 µg mL^−1^ kanamycin for plasmid maintenance on agar plates and in liquid culture.

All experiments in liquid culture were inoculated in triplicate to an optical density at 750 nm (OD) of 0.5 in 50 mL BG-11_H_ supplemented with 50 µg mL^−1^ kanamycin. Cultures were induced with 0.1 mg mL^-1^ L(+)-rhamnose monohydrate (Carl-Roth) after 24 h of growth. Samples were harvested every 24 h to measure the OD and extract pigments.

The production experiment in high CO_2_ conditions was carried out in a HDC 6.10 cultivation setup (CellDeg) in P4-TES CPH medium as previously described (Russo et al., 2022) with small modifications: the cultures were supplemented with 50 µg mL^−1^ kanamycin and induced with 1 mg mL^-1^ of rhamnose after 24 h of growth. HDC culture vessels were agitated by shaking at 150 rpm.

### 2.2. Construction of plasmids

All cloning was performed in *Escherichia coli* NEB 5-alpha (New England Biolabs) for plasmid propagation. Plasmids were generated by GoldenGate cloning using the CyanoGate syntax (Vasudevan et al., 2019). The self-replicating plasmid pPMQAK1-T (Addgene kit #1000000146, gift from Alistair McCormick) was used for final construct assembly in Level T. For domestication (removal of internal BbsI recognition sites in *BvDODA1* gene) and generation of individual CyanoGate compatible parts, DNA sequences were either amplified from existing plasmids or custom-synthesized as double stranded DNA fragments (gBlock, Integrated DNA technologies). BbsI recognition sites were added for Level 0 part generation by PCR with Q5^®^ Hot start High-Fidelity DNA Polymerase (New England Biolabs). Primers sequences used for individual part generation are shown in Table S1 (Supplementary Material). GoldenGate assemblies were performed using a standard protocol (Weber et al., 2011) with small modifications: assembly reactions were incubated in a thermocycler with 30 cycles of restriction enzyme activation (BbsI-HF or BsaI-HFv2, New England Biolabs) at 37 °C for 5 min followed by activation of the T4 DNA ligase for 5 min at 16°C. A final restriction enzyme inactivation step was added at 85 °C for 20 min. Level 0 and final Level T constructs were verified using Sanger sequencing (Eurofins Genomics) and colony PCR. Plasmid maps of generated level T shuttle vectors (Table 1) are deposited on Zenodo with the DOI:10.5281/zenodo.15673972.

**Table 1.** *Synechocystis* sp. PCC 6803 and *S. elongatus* PCC 7942 strains generated in this study.

| Strain ID | Plasmid | Description |
| --- | --- | --- |
| Syn-N<sup>Kan</sup> | pPMQAK1-T<br>(Vasudevan et al., 2019) | *Synechocystis* sp. PCC 6803 carrying self-replicative shuttle plasmid (CyanoGate Kit), (Vector ID pCAT.000); no gene of interest, selection markers: Ampicillin, Kanamycin. |
| Syn-Bx | pSH023 | *Synechocystis* sp. PCC 6803 carrying plasmid with promoter P<sub>RhaBAD</sub>-Bx operon (*CYP76AD1-HA*, *BvDODA1-HA*)- T<sub>cpc_operon</sub>, P<sub>J23116</sub>-*rhaS*)- T<sub>cpc_operon</sub>; backbone: pPMQAK1-T. |
| Syn-Bx<sup>AroG*</sup> | pSH018 | *Synechocystis* sp. PCC 6803 carrying plasmid with promoter P<sub>RhaBAD</sub>-Bx operon (*CYP76AD1-HA*; *BvDODA1-HA*)-T<sub>cpc_operon</sub>, promoter P<sub>J23116</sub>-*aroG\**-*his*, *rhaS*-T<sub>cpc_operon</sub>; backbone: pPMQAK1-T. |
| Se-N<sup>Kan</sup> | pPMQAK1-T<br>(Vasudevan et al., 2019) | *S. elongatus* PCC 7942 carrying pPMQAK1-T. |
| Se-Bx<sup>AroG*</sup> | pSH018 | *S. elongatus* PCC 7942 carrying plasmid with promoter P<sub>RhaBAD</sub>-Bx operon (*CYP76AD1-HA*; *BvDODA1-HA*)-T<sub>cpc_operon</sub>, promoter P<sub>J23116</sub>-*aroG\**-*his*, *rhaS*-T<sub>cpc_operon</sub>; backbone: pPMQAK1-T. |

### 2.3. Generation of cyanobacterial strains

Plasmids were transferred into cyanobacterial recipient strains by triparental mating as described previously (Zedler et al., 2023) and selected on agar plates with 50 μg mL^−1^ kanamycin. The presence of target plasmids in colonies was verified by colony PCR. DNA from colonies was extracted by boiling of a single colony resuspended in 50 µL of ddH_2_O at 95°C for 10 min. Subsequently, 2 µL of the DNA extract were used in a PCR reaction with primers specified in Table S1. An overview of all generated strains from this study is shown in Table 1.

### 2.4. Preparation of whole cell lysates and determination of protein content

Whole cell lysates were prepared as previously described (Russo et al., 2019b) with the exception that an equivalent of OD = 10 was harvested after 9 days of cultivation and lysed in 0.5 mL of lysis buffer. The protein content of cleared cell lysates was estimated with a Pierce bicinchoninic acid protein assay kit (Thermo Fisher Scientific) in a 96-well plate format.

### 2.5. SDS-PAGE and immunoblotting

An equivalent of 10 µg total protein extracted from cleared cell lysates was denatured at 50 °C for 10 min in sample loading buffer (containing 2% SDS and 0.1 M dithiothreitol). Proteins were subsequently separated on 4–12% Criterion™ XT Bis-Tris Protein Gel (Bio-Rad Laboratories) using 3-(N-morpholino) propanesulfonic acid (MOPS) as the running buffer at 100 V. Proteins were then transferred onto a 0.45-μm pore size nitrocellulose membrane using a Trans-Blot® Turbo transfer system (Bio-Rad Laboratories) for immunoblotting. Membranes were blocked with 5% skimmed milk powder (w/v) in TBS-T buffer (10 mM Tris-HCl (pH 8.0), 150 mM NaCl, and 0.05% Tween 20) for 60 min at room temperature. The nitrocellulose membrane was either incubated with an anti-HA antibody produced in rabbit (dilution 1:5000, Sigma-Aldrich, product no.: H6908) or an anti-His antibody produced in mouse (dilution 1:1000, Novagen, product no.: 70796) overnight at 4 °C. Membranes were washed three times with TBS-T and incubated for 60 min at room temperature with an anti-rabbit or anti-mouse HRP-conjugated secondary antibody (Promega) at a 1:5000 dilution. After three more wash steps in TBS-T, the chemiluminescent signal was developed using a Clarity Max Western ECL Substrate (Bio-Rad Laboratories). Chemiluminescence was detected using a ChemiDoc imager (Bio-Rad Laboratories).

### 2.6. Fluorescence measurements in culture supernatants

The supernatant from cultures was harvested by centrifugation at 9,300 x g for 5 min at 4 °C. A volume of 100 µL culture supernatant was transferred to black 96-well plates. Fluorescence measurements were recorded in a VANTAstar® plate reader (BMG Labtech) with the excitation set to 475 nm and emission measured at 515 nm. A bandwidth of 15 nm (excitation and emission) with a focal height of 5 mm and a fixed gain of 1000 was used. Significance was tested with a multiple unpaired Student’s t-test and the false discovery rate (FDR) was controlled by the Benjamini-Krieger-Yekutieli two-stage procedure.

### 2.7. L-DOPA feeding assay

For L-3,4-Dihydroxyphenylalanine (L-DOPA, Sigma-Aldrich) feeding experiments, two sets of cultures in triplicate were grown and induced with 1 mg mL^-1^ rhamnose after 24 h. After 48 h, L-DOPA was added to one set of cultures at a final concentration of 100 µM. As a control, the equivalent volume of fresh medium was added to the other set. An L-DOPA stock solution (10 mM L-DOPA and 0.6% w/v ascorbic acid) was prepared fresh for each experiment. Significance was tested with a multiple unpaired Student’s t-test and the false discovery rate (FDR) was controlled by the Benjamini-Krieger-Yekutieli two-stage procedure.

### 2.8. Pigment analysis

Chlorophyll *a* and carotenoid content of samples was determined by methanol extraction from frozen pellets as previously described (Russo et al., 2019b). Pigment content in mg L^-1^ was calculated for chlorophyll *a* (Ritchie, 2006) and total carotenoids (Myers et al., 1980).

### 2.9. Semi-synthesis of a phenylalanine-betaxanthin standard

A semi-synthetic phenylalanine-betaxanthin standard was prepared in-house following previously established method (Cabanes et al., 2014) with several modifications. Two grams of commercial *Beta vulgaris* (beet) powder was dissolved in 20 mL ddH_2_O. The mixture was then filtered with a 125 µM E-D-SCHNELLSIEB^®^ nylon filter to remove remaining insoluble debris. Subsequently, 2 g of Lewatit VP OC 1065 (Sigma-Aldrich) were added to the filtrate and the pH titrated to pH 11.4 with aqueous ammonia. After a 10 min incubation step with constant stirring at room temperature, the pH was reduced to 5.0 with glacial acetic acid. The mixture was incubated for 30 min with constant stirring. Lewatit beads were then collected on a filter and thoroughly rinsed first with ddH_2_O followed by methanol. The beads were then carefully dried under a nitrogen gas stream. Beads were resuspended in aqueous ammonia and incubated for 30 min at pH 11.4.

Following the addition of 500 mg phenylalanine, the pH of the solution was adjusted to 5.0 with glacial acetic acid. The condensation product was stirred for another 30 min before storage at −20 °C until further use.

### 2.10. Liquid chromatography mass spectrometry (LC-MS)

For LC-MS analysis, the culture medium was harvested by centrifugation at 9,300 x g, 5 min at 4 °C and samples were stored at −80 °C until analysis. Ultra-high-performance LC coupled with high resolution MS was carried out using a Vanquish UHPLC system (Thermo Fisher Scientific) with a VF-P10-A binary pump and a VF-A10-A auto sampler which was set to 10 °C and equipped with a 25 µL injection syringe and a 100 µL sample loop. The column, Accucore® C-18 RP (100 × 2.1 mm; 2.6 µm), was kept at 25 °C within the column compartment VH-C10-A. Eluent A was water with 2% acetonitrile and 0.1% formic acid. Eluent B was pure acetonitrile. All solvents and water were LC-MS grade. The gradient for sample separation started with 0% B to 2.2 min, followed by an increase to 11% B over 10 min, and 90% B over a further 4 min. It was then kept at 90% B for 2 min and re-equilibrated to 0% B over 1 min. Flow rate was kept constant at 0.5 mL min^-1^. The Vanquish UHPLC system was also equipped with a variable wavelength detector (VC-D40-A-01). UV-Vis data was collected from 190 to 680 nm with a bunchwidth of 1 nm. Mass spectra were recorded with an Orbitrap Exploris 480 MS (Thermo Fisher Scientific) coupled to a heated electrospray (HESI) source. For monitoring two full scan modes were selected with the following parameters. Polarity: positive; scan range (*m*/*z*): 100 to 1500; resolution: 180,000; AGC target: standard; maximum injection time mode: auto. General settings: sheath gas flow rate (arb): 50; auxiliary gas flow rate (arb): 10; sweep gas flow rate (arb): 1; spray voltage (kV): 3.5; capillary temperature (°C): 325; RF lens (%): 50; vaporizer temperature (°C): 350; acquisition time (min): 0.7 - 16.2. Data analysis and peak area calculation were done on Freestyle 1.8 SP2 (Thermo Fisher Scientific).

## 3. Results

### 3.1. Precursor availability limits betaxanthin production in Synechocystis sp. PCC 6803

In *Beta vulgaris*, two enzymes are required for the biosynthesis of Bx: a cytochrome P450 (CYP76AD1, UniProtKB: I3PFJ5), that converts tyrosine into L-DOPA, and a L-DOPA dioxygenase (BvDODA1, UniProtKB: I3PFJ9), which converts L-DOPA into betalamic acid. Betalamic acid can then spontaneously condense with amines and amino acids to form Bx (Fig. 1A). To establish a Bx pathway in cyanobacteria, genes encoding C-terminally HA-tagged variants of CYP76AD1 (57.1 kDa) and BvDODA1 (32.2 kDa) were expressed in Syn6803 (hereafter, Syn-Bx) under the control of the rhamnose-inducible *rhaBAD* promoter. For prhaBAD transcriptional activation, the regulator RhaS was co-expressed under the control of the constitutive low-to-medium strength J23116 promoter (Fig. 1B, Table 1). Successful transfer of the self-replicative plasmid was confirmed by colony PCR (Fig. S1). To determine whether the two proteins were produced, cultures of Syn-Bx were induced with rhamnose and grown for immunoblot analysis. The presence of CYP76AD1 and BvDODA1 in Syn-Bx cleared cell lysates was confirmed at the predicted molecular weight of 32 kDa and 57 kDa, respectively (Fig. 1C). In microbial production systems, betaxanthins are known to accumulate in the medium (Guerrero-Rubio et al., 2019). Therefore, medium samples were harvested from cultures every 24 h to determine if betaxanthins were produced in Syn-Bx. Taking advantage of the intrinsic betaxanthin fluorescence, we conducted a fluorescence analysis using previously determined excitation and emission maxima for betaxanthins (Gandía-Herrero et al., 2005a). This did not provide any indication of betaxanthin accumulation (Fig. 1D). Overall, our results showed that CYP76AD1 and BvDODA1 were expressed in Syn-Bx, however no detectable amount of betaxanthin could be measured.

**Fig. 1.**
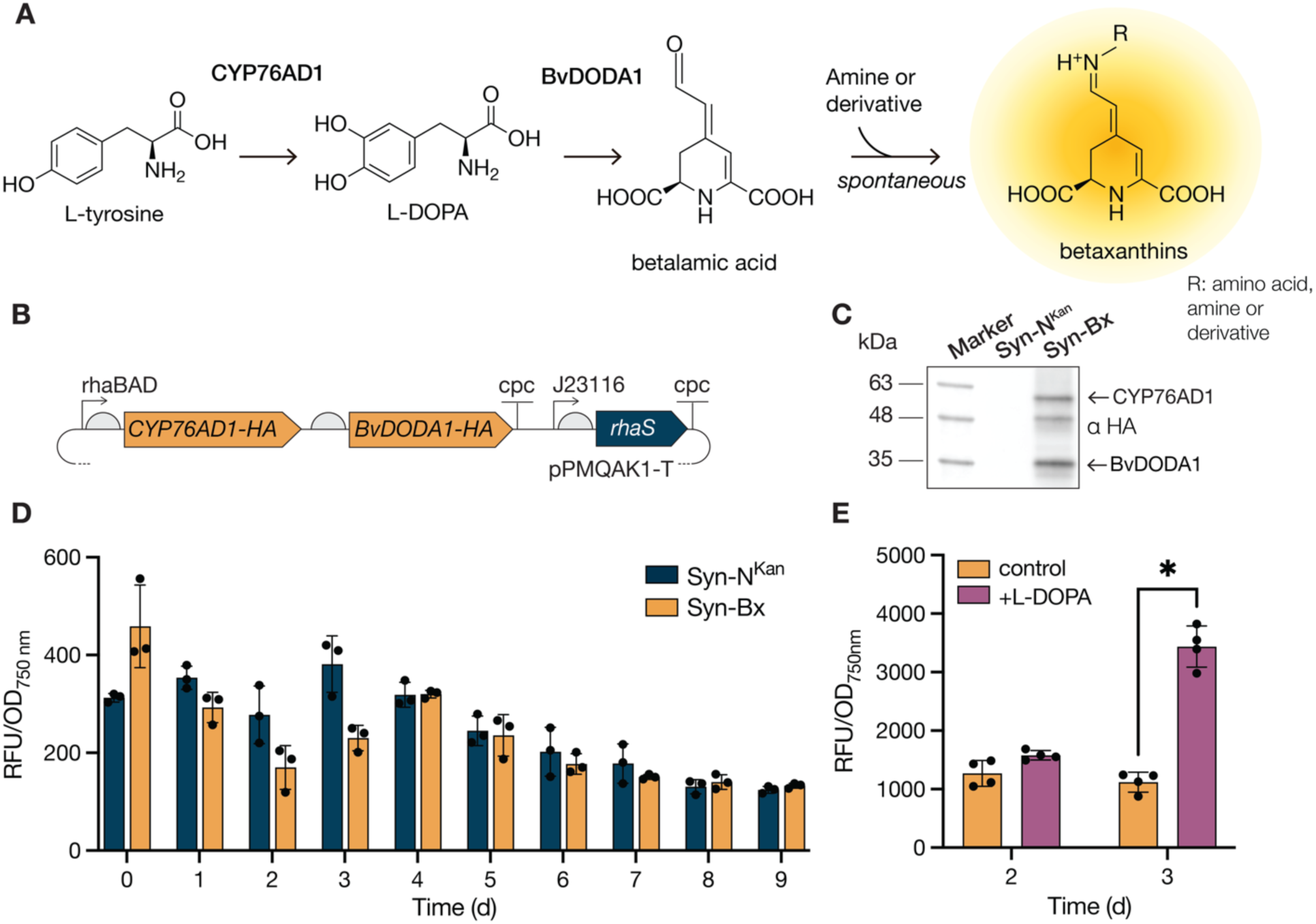
Engineering of a minimal pathway for betaxanthin (Bx) biosynthesis in Syn6803. **A** Overview of Bx pathway in *Beta vulgaris*. **B** Construct design of extrachromosomal expression of a Bx operon (*CYP76AD1-HA* and *BvDODA1-HA)* under the control of the rhaBAD promoter. The transcription factor-encoding gene *rhaS* was expressed from the same vector with a constitutive J23116 promoter. **C** Immunodetection of CYP76AD1 and BvDODA1 in cleared cell lysates with an Anti-HA antibody (αHA). Syn-Bx denotes the strain engineered for Bx biosynthesis, Syn-N^Kan^ is a negative control strain carrying the plasmid pPMQAK1-T. **D** OD-normalized fluorescence (Ex: 475/15, Em: 515/15) of culture medium measured over 9 days of cultivation. Cultures were induced at day 1. Bars show averages (n = 3), error bars: ± SD. **E** OD-normalized fluorescence (Ex: 475/15, Em: 515/15) of culture medium with and without the addition of L-DOPA after 2 days of growth. Bars show averages (n = 4), error bars: ± SD.

To troubleshoot the lack of betaxanthin production, we started by investigating whether the first step of the pathway, namely the conversion of tyrosine to L-DOPA by CYP76AD1, was the bottleneck. To test this, a Syn-Bx culture was induced, grown for 2 days, and then fed with 100 µM L-DOPA. We observed that normalized fluorescence was approximately three times higher in L-DOPA-fed cultures when compared to their counterparts where no L-DOPA was added (Fig. 1E). To identify the main betaxanthin responsible for the increased fluorescence levels, the culture medium was analyzed by liquid chromatography mass spectrometry (LC-MS). Despite a thorough search using the monoisotopic masses of betalamic acid and all 20 amino acid betaxanthins (Esteves et al., 2022), neither betalamic acid nor betaxanthins were identified. One possible explanation is that betalamic acid spontaneously condensed with multiple, readily available, amino acids or amines and no dominant product emerged. Thus, complicating the identification of the specific betaxanthins formed

### 3.2. Overproduction of tyrosine and phenylalanine leads to the accumulation of phenylalanine-betaxanthin

Our results suggested that the betaxanthin pathway was functional in Syn6803 but limited by L-DOPA availability. Given that amount of CYP76AD1 protein was observed at similar levels to BvDODA1 (Fig. 1C), we hypothesized that the L-DOPA limitation could derive from insufficient tyrosine supply to CYP76AD1 (Fig. 1A). To test this hypothesis, a feedback-resistant *aroG* (*aroG^fbr^*) from *E. coli* was added in an operon with the transcription factor *rhaS* to the Bx plasmid (Fig. 2A, Table 1). AroG is a 3-deoxy-D-arabino-heptulosonate-7-phosphate (DAHP) synthase that regulates one of the first steps of the shikimate pathway. While AroG is typically inhibited by phenylalanine, in AroG^fbr^ the amino acid residues responsible for the feedback inhibition have been mutated. In a previous study, expression of *aroG^fbr^* in Syn6803 led to 100 times increase in phenylalanine concentrations, and 10 times increase in tyrosine and tryptophan concentrations (Brey et al., 2020). The generated plasmid was conjugated into Syn6803 to generate the strain Syn-Bx^AroG*^ (Fig 2). Successful strain generation was validated by colony PCR (Fig. S1). Cultures of the Syn-Bx^AroG*^ strain were then grown for immunoblot and product analysis. Immunoblot analysis of the Syn-Bx^AroG*^ cleared cell lysates confirmed the presence of all three proteins CYP76AD1, BvDODA1 and AroG^fbr^ (Fig. 2B). Growth analysis showed that, in comparison to the negative control strain and the Syn-Bx strain, Syn-Bx^AroG*^ entered stationary phase earlier and achieved an approximately 30% lower final OD (Fig. S2A). This was supported by the pigment analysis which showed that the chlorophyll concentrations of Syn-Bx^AroG*^ were approximately 25-35% lower in comparison to the other two strains (Fig. S2B) throughout most of the experiment. On the other hand, carotenoid concentrations were similar among all strains (Fig. S2C).

**Fig. 2.**
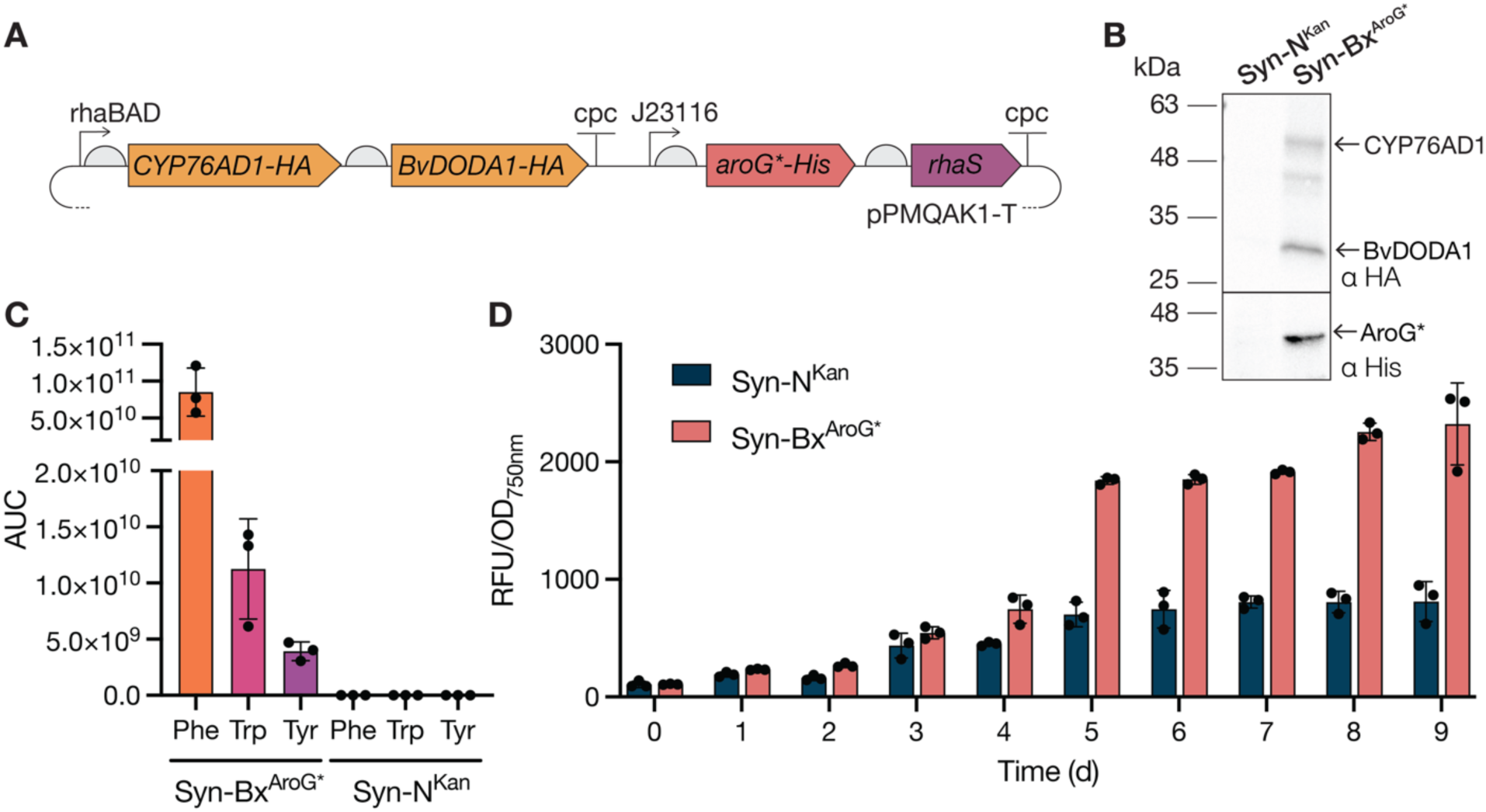
Co-expression of *aroG^fbr^* increases the aromatic amino acid pool in a betaxanthin producing Syn6803 strain. **A** Construct design for co-expression of a Bx operon in conjunction with a pJ12116-controlled operon of a feedback-resistant *aroG\**. Syn-Bx^AroG*^ denotes the strain engineered for Bx biosynthesis with AroG^fbr^, Syn-N^Kan^ is a negative control strain carrying the plasmid pPMQAK1-T. **B** Immunodetection of CYP76AD1 and BvDODA1 with an Anti-HA antibody (αHA) and AroG*-His detection with an Anti-His antibody (αHis) in cleared cell lysates. **C** LC-MS evidence of amino acids phenylalanine (Phe), tryptophan (Trp) and tyrosine (Tyr) in the culture medium. **D** OD-normalized fluorescence (Ex: 475/15, Em: 515/15) of culture medium measured over 9 days of cultivation. Cultures were induced at day 1. Bars show averages (n = 3), error bars: ± SD.

Given that the presence of AroG^fbr^ should increase the amount of phenylalanine, tyrosine, and tryptophan, we searched for chromatographic signals of these three amino acids in the culture medium of Syn-Bx^AroG*^. An excess of phenylalanine, tyrosine, and tryptophan could be detected, thus suggesting that AroG^fbr^ was functioning as expected (Fig. 2C). This was also reflected in a subsequent fluorescence analysis. Normalized fluorescence of the Syn-Bx^AroG*^ culture medium was significantly higher than that of Syn-N^Kan^ from day 4 (p = 0.01) until the end of the experiment (p < 0.001), suggesting the formation of betaxanthins (Fig. 2D). Normalized fluorescence peaked at day 5 and gradually decreased until day 9 (Fig. 2D). Therefore, culture medium harvested at days 5 and 9 was further analyzed by LC-MS.

We searched for chromatographic signals of the respective betaxanthins derived from condensation of betalamic acid with the overproduced aromatic amino acids (Phe, Trp, Tyr) in the Syn-Bx^AroG*^ culture medium. However, only the phenylalanine-betaxanthin (Phe-Bx) could be putatively identified. The identity of Phe-Bx was confirmed by comparing the associated retention time and mass with those of a semi-synthetic standard (Fig. 3). The two peaks observed in the reference standard likely correspond to the 2*S/S* and 2*S/R* Phe-Bx isomers (Fig. 3, bottom). As observed in previous studies, only the 2*S/S* isomer is produced in vivo (Fig. 3, center) (Gandía-Herrero et al., 2005b). A standard curve was then measured (Fig. S4) and used to determine that 0.11 ± 0.01 mg L^-1^ and 0.08 ± 0.03 mg L^-1^ of Phe-Bx were produced after 5 and 9 days of cultivation, respectively.

**Fig. 3.**
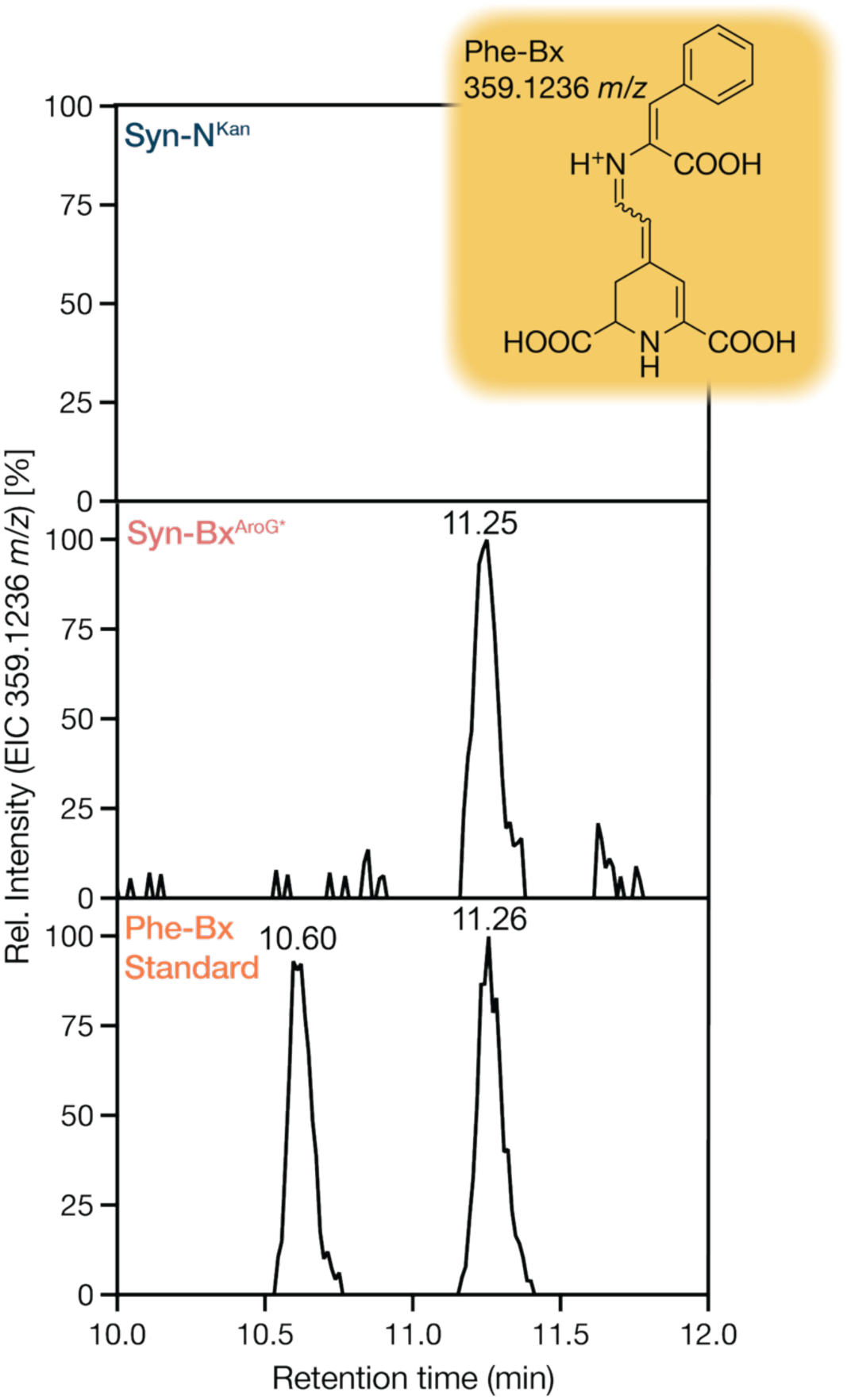
LC-MS evidence of phenylalanine-betaxanthin (Phe-Bx) production in the Syn-Bx^AroG*^ strain. Extracted ion chromatograms (EIC) of the negative control strain Syn-N^Kan^ (top), Syn-Bx^AroG*^ (center) and 5 µM of the Phe-Bx semi-synthetic standard (bottom). The structural formula of Phe-Bx and its associated monoisotopic mass in positive ion mode are shown in the inset.

Although betaxanthins were successfully produced in Syn6803, the yield was relatively low. To determine whether a different cyanobacterial chassis would be more suitable for Bx biosynthesis, we tested the Bx operon combined with AroG^fbr^ in *Synehococcus elongatus* PCC 7942 (hereafter Syn7942). The pSH018 plasmid (Table 1) was conjugated into the wildtype strain (Table 1) to generate the strain Se-Bx^AroG*^ (Fig. S1). Immunoblot analysis of induced cultures showed that BvDODA1 was produced to high levels whereas only a weak signal was seen for CYP76AD1 (Fig. S3A) and AroG^fbr^ was not detected by immunoblotting (data not shown).Subsequent fluorescence analysis of the medium also showed no major differences between a negative control strain (Se-N^Kan^) and the Se-Bx^AroG*^ strain (data not shown). Given the low levels of CYP76AD1, and the absence of AroG^fbr^, we hypothesized that the bottleneck was the supply of L-DOPA to BvDODA1. To test this hypothesis, we performed an L-DOPA feeding experiment. Cultures of Se-Bx^AroG*^ were grown, induced, and after 2 days, L-DOPA was added to a concentration of 100 µM. Analysis of the culture medium 24 h after L-DOPA addition showed a six times higher normalized fluorescence signal in L-DOPA-fed cultures in comparison to cultures where L-DOPA was not added (Fig. S3C). An LC-MS search in culture medium samples using the monoisotopic masses of betalamic acid and all 20 amino acid betaxanthins did not identify any betaxanthin precursors or products. This was a similar result to what was observed in the Syn-Bx feeding experiment (Fig. 1E). Overall, these results suggested that the low levels of CYP76AD1, together with the absence of AroG^fbr^, was the cause behind the non-functional betaxanthin pathway in Se-Bx^AroG*^.

### 3.3. High-density cultivation significantly improves betaxanthin yields in Syn-Bx^AroG^*

The expression of *aroG^fbr^* had a negative impact on growth and led to an early entrance into stationary phase in Syn-Bx^AroG*^ (Fig. S2C). This is likely due to the metabolic burden of overproducing multiple amino acids (Brey et al., 2020). Thus, we decided to test whether optimization of the growth conditions could further boost Phe-Bx titers in Syn6803. To this end, we tested cultivation of Syn-Bx^AroG*^ in a nutrient-rich medium with CO_2_-enriched air. After induction, the OD and fluorescence of the culture medium were monitored daily. A significant increase in the Syn-Bx^AroG^* culture medium fluorescence was observed starting 4 days after induction (p = 0.01). It then continuously increased until day 9 (Fig. 4A). Based on LC-MS analysis, the volumetric productivity of Phe-Bx was 2.4 mg/L and 15.3 mg/L at days 5 and 9, respectively (Fig. 4B). A titer approximately 150 times higher than what was achieved in standard growth conditions.

**Fig. 4.**
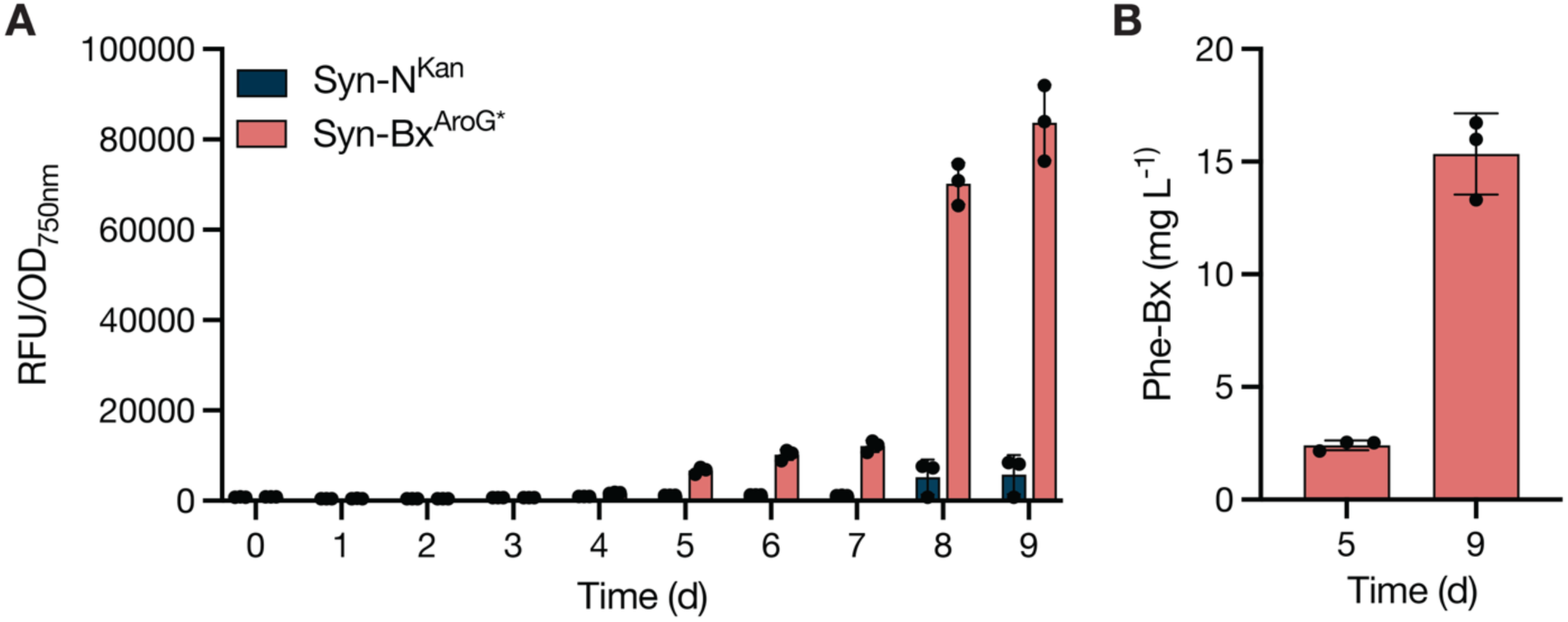
Betaxanthin production in Syn-Bx^AroG*^ strain in high density cultivation. **A** OD-normalized fluorescence (Ex: 475/15, Em: 515/15) of culture medium measured over 9 days of cultivation with nutrient rich P4-TES CPH medium and high CO2 conditions. Cultures were induced at day 1. Bars show averages (n = 3), error bars: ± SD. **B** Phe-Bx titers determined by LC-MS.

## 4. Discussion

In this study we used a beetroot cytochrome P450, CYP76AD1, and L-DOPA dioxygenase, BvDODA1, to establish betaxanthin pigment biosynthesis in cyanobacteria. Introduction of the feedback-resistant *aroG^fbr^* together with the betaxanthin pathway led to the production of 0.11 mg L^-1^ of Phe-Bx. Cultivation in high CO_2_ conditions then increased titers almost 150 times and allowed for the fully photoautotrophic production of of Phe-Bx at a titer of 15.3 mg L^-1^.

When comparing to other efforts to produce Phe-Bx, this value is higher than the 11 mg L^-1^ achieved with a Phe-fed *E. coli* strain expressing a 4,5-DOPA-extradiol-dioxygenase from *Gluconacetobacter diazotrophicus* (Guerrero-Rubio et al., 2019). Our 15.3 mg L^-1^ Phe-Bx titer is also in the range of what has been achieved for betanin production in glucose-fed *Saccharomyces cerevisiae* (16.8 – 30.8 mg L^-1^ (Babaei et al., 2023; Grewal et al., 2018; Zhang et al., 2023) The highest titer of betalains achieved to date was 22.9 g L^-1^ of dopaxanthin (i.e., condensation production of L-DOPA and betalamic acid) using an *E. coli* strain with substantial metabolic adjustments of the shikimate pathway and two gene copies of an evolved DODA (Jiang et al., 2024).

In this work, the addition of the feedback-resistant *aroG^fbr^*was crucial for high-titer betaxanthin production. This was likely due to the larger Tyr pool available for CYP76AD1-catalyzed production of L-DOPA. Expression of *aroG^fbr^* also led to the overaccumulation of Phe, Trp and Tyr in the medium (Fig. 2C). However, only Phe-Bx was detected. This can be attributed to (i) the accumulation of significantly more Phe than Trp and Tyr, and (ii) the increased solubility of Phe (approximately twice that of Trp and 50 times higher than Tyr) (Fleck and Petrosyan, 2014). Residual L-DOPA or betalamic acid were not detected in any of the betaxanthin-producing strains. This suggests that Tyr supply is still the rate-limiting step of the reaction. Therefore, additional manipulation of the shikimate pathway, such as the introduction of the feedback-resistant TyrA^fbr^, could further increase betaxanthin titers (Brey et al., 2020). However, given that manipulation of the shikimate pathway is a universal strategy for betalain overproduction (Khan and Polturak, 2025), it may be difficult to produce amino acid betaxanthins in cyanobacteria that do not contain Phe or Trp. One potential strategy could be a two-partner co-cultivation approach: one producing, and secreting, betalamic acid and the second partner secreting an overproduced amino acid for condensation. This was successfully implemented with two *E. coli* strains in co-culture and led to the production of 287.69 mg L^-1^ of histidine-betaxanthin (Hou et al., 2020).

An alternative that does not rely on amino acid supply for the condensation reaction is to produce the more stable red-violet betacyanins (Esteves et al., 2022). This can be achieved by introduction of UDP-glycosyltransferases that can glycosylate the unstable betanidin (product of the condensation of betalamic acid with cyclo-DOPA) to form betanin. Alternatively, cyclo-DOPA can first be glycosylated by a UDP-glycosyltransferases to form its glucoside, which then condenses with betalamic acid to form betanin (Glitz et al., 2025). Given that the CYP76AD1 in our system can produce cyclo-DOPA (Timoneda et al., 2019), betanin production should require only the introduction of appropriate UDP-glycosyltransferases.

Due to their inherent capacity to produce heme-containing proteins, cyanobacteria have become choice organisms for the production of cytochrome P450s (Russo et al., 2019a). There is the added benefit that membrane-bound plant cytochrome P450s typically get inserted into the cyanobacteria thylakoid membranes, thus allowing for NADPH-independent light-driven catalysis (Wlodarczyk et al., 2016). If this is also the case with the CYP76AD1 used in this study, then previous synthetic biology strategies to improve cytochrome P450 production, stability, and catalysis could also be applied to betalain biosynthesis. These include improving thylakoid membrane targeting (Hanamghar et al., 2025), fusion of the cytochrome P450 to dedicated electron donors (Mellor et al., 2019; Sutardja et al., 2024), and elimination of competing electron sinks (Spasic et al., 2022). It is worth noting that the strategy applied to produce betaxanthins in Syn6803 was not successful in Syn7942. This was likely due to low levels or potential absence of CYP76AD1 and AroG^fbr^ (Fig. S3). The low level/absence of AroG^fbr^ could be explained by the difference in strength (i.e., weaker in Syn7942) of the constitutive promoter J23116. This is supported by a recent study that successfully expressed *aroG^fbr^* in Syn7942 when under the control of the trc promoter. The strong expression of *aroG^fbr^*, when coupled with the overexpression of the endogenous shikimate kinase gene *aroK*, significantly increased the carbon flow into the shikimate pathway (Usai et al., 2022). This approach, together with potential cytochrome P450 improvements described above, could unlock betaxanthin biosynthesis in Syn7942.

In conclusion, our work showcases the biosynthetic potential of cytochrome P450-containing pathways in cyanobacteria and opens the door to sustainable production of plant-derived betalain pigments in a microbial host.

## Acknowledgements

This work was supported by a PhD scholarship from the Federal State of Thuringia granted to SSH, and the Deutsche Forschungsgemeinschaft (DFG, German Research Foundation) SFB 1127 ChemBioSys, project number 239748522 (JAZZ, DAR). The Exploris 480 mass spectrometer was funded by the state of Thuringia (2023 INST 275/535-1 FUGG) and by the Deutsche Forschungsgemeinschaft (DFG, German Research Foundation) 529714284 with means of the 91bGG proposal.

## Author contributions (CRediT)

**SSH**: Conceptualization, Methodology, Investigation, Formal Analysis, Investigation, Data Curation, Writing - Original Draft, Writing – Review & Editing, Visualization, Funding Acquisition. **DAR**: Conceptualization, Methodology, Investigation, Formal Analysis, Investigation, Data Curation, Writing – Original Draft, Writing – Review & Editing, Visualization. **SBM:** Conceptualization, Methodology, Formal Analysis, Writing – Review & Editing. **JAZZ**: Conceptualization, Methodology, Resources, Formal Analysis, Writing - Original Draft, Writing – Review & Editing, Visualization, Funding Acquisition.

