## Supplementary Material for "Establishing betaxanthin pigment biosynthesis in cyanobacteria"

### Overview Supplementary Material

**Fig. S1** Colony PCR of generated cyanobacterial strains.

**Fig. S2** Growth and pigment analysis of Syn-Bx and Syn-Bx$^{AroG*}$ strains

**Fig. S3** Protein and Bx detection in Se-Bx$^{AroG*}$

**Fig. S4** LC-MS calibration curve of synthesized Phe-Bx standard.

**Table S1** Primers used in this study.

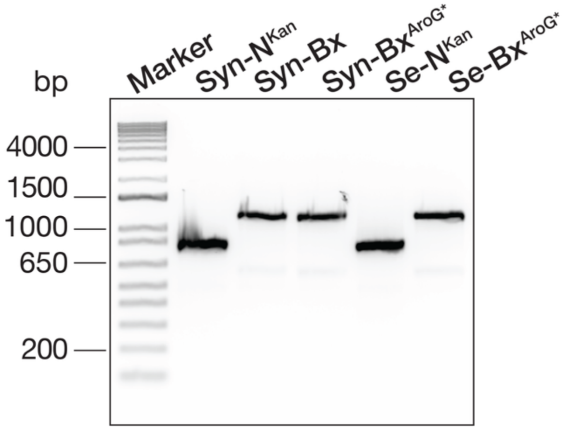


**Fig. S1**. Confirmation of plasmid presence in *Synechocystis* sp. PCC 6803 (Syn) and *Synechococcus elongatus* PCC 7942 (Se) strains generated for this study by PCR-amplification of plasmid fragments from single colonies. The primer pair CDS1_SeqF and CDS_SeqR was used for strains with plasmids without a gene of interest transformed with pPMQAK1-T (Syn-$N^{Kan}$, Se-$N^{Kan}$). The primer pair CDS1_SeqF and Mid_CYP76_SeqR was used for all other strains containing the Bx operon. In the Syn-Bx and Syn-$Bx^{AroG*}$ strains the plasmids pSH023 and pSH018 were inserted, respectively.

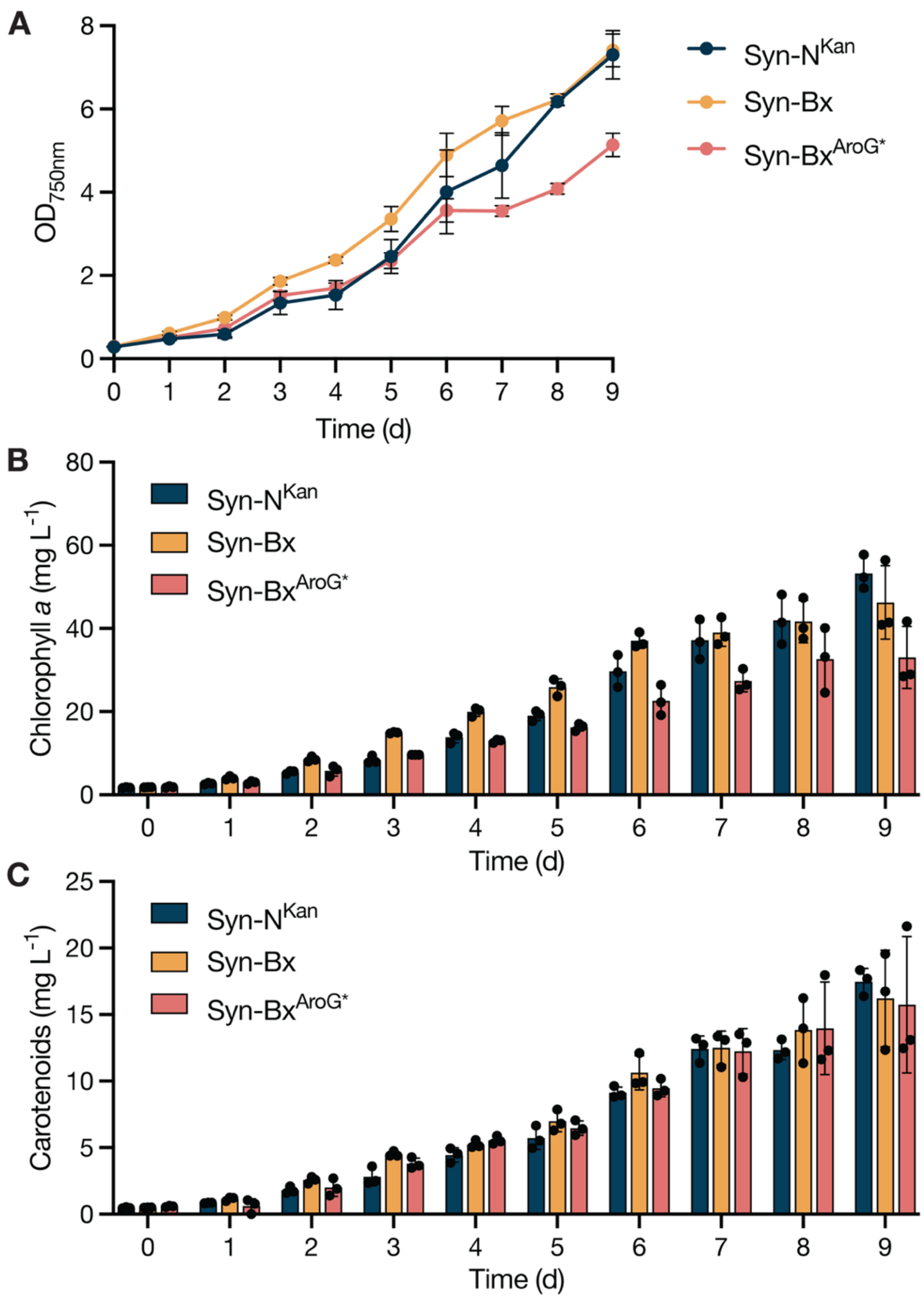


**Fig. S2**. Growth and pigment analysis of *Synechocystis* sp. PCC 6803 strains. Syn-N$^{Kan}$: negative control strain, Syn-Bx: strain producing CYP76AD1 and BvDODA1. Syn-Bx$^{AroG*}$: strain producing CYP76AD1, BvDODA1 and AroG*. **A** Growth curves recorded over 9 days of cultivation at 30°C with 50 µmol photons m$^{-2}$ s$^{-1}$ in BG-11$_H$ medium supplemented with 50 µg mL kanamycin. Cultures were induced after 1 day with 1 mg mL$^{-1}$ rhamnose. **B** Estimation of chlorophyll *a* content in mg L$^{-1}$. **C** Estimation of carotenoid content in mg L$^{-1}$. n = 3, error bars: ± SD.

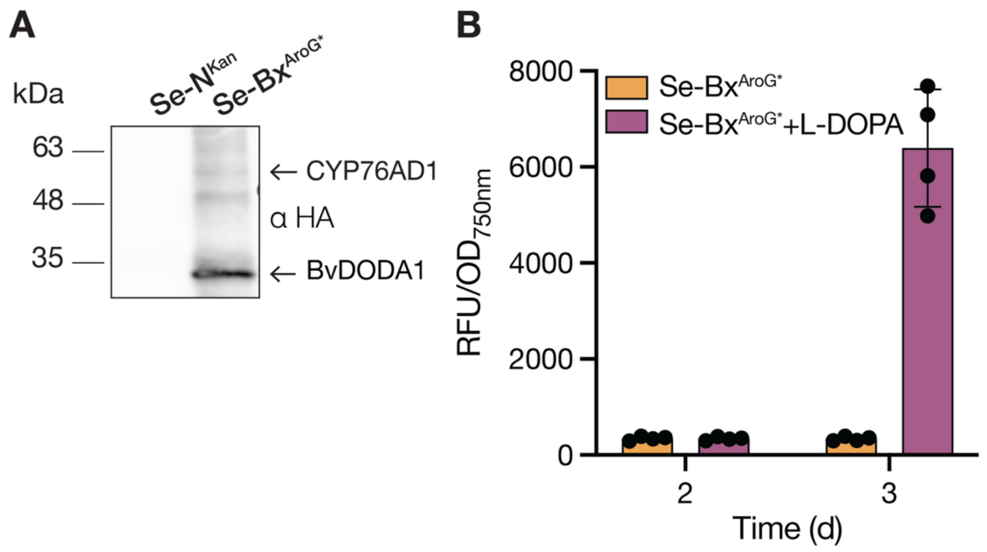


**Fig. S3**. Engineering *Synechococcus elongatus* PCC 7942 to produce betaxanthins. **A** Immunodetection of CYP76AD1 and BvDODA1 with an anti-HA antibody (αHA) in cleared lysates. AroG*-His detection with an anti-His antibody (αHis) did not show any detectable signal (data not shown). **B** OD-normalized fluorescence (Ex: 475/15, Em: 515/15) of Se-Bx$^{AroG*}$ detected in culture medium with and without the addition of L-DOPA to a final concentration of 100 μM. Addition was done on day 2. Bars show averages (n = 4), error bars: ± SD.

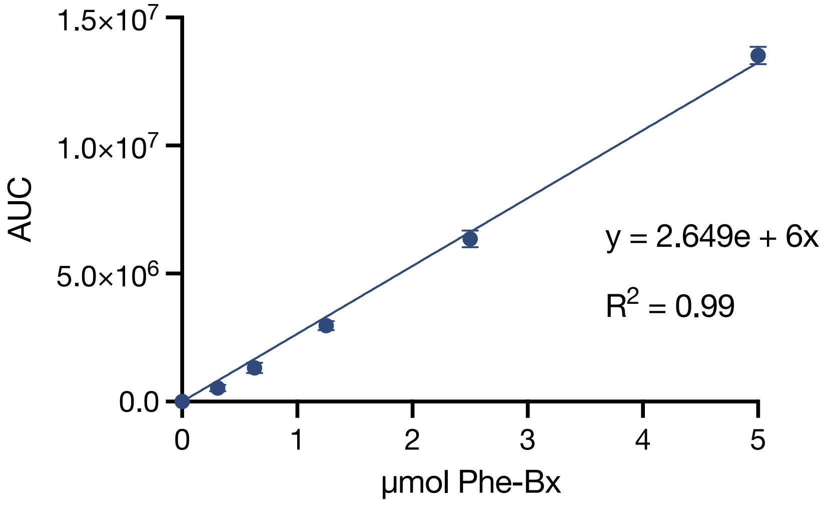


**Fig. S4.** Linear regression function for the LC-MS data of the phenylalanine-betaxanthin (Phe-Bx) standard. The peak area of the analyte was plotted against the concentration of the calibration standard (n = 3). The error bars represent the mean ± SD.

**Table S1.** List of primers used in this study.

| Primer name | Sequence 5'- 3' | Application of primer |
|---|---|---|
| BbsI_CYP76-HA_GG_F | CGGCGAAGACAGAATGAAATGGATCACGCAACATTAGCAATG | Generation of Level 0 part *CYP76AD1-HA* |
| BbsI_CYP76-HA_GG_R | ATGCGAAGACAAAAGCTCATGCGTAGTCCGGGACGTCGTACGGATAATACCTAG | |
| Silentmut_BvDODA1_F | ACACATGGAAACCCAATATTAACAGTAGAAGATACACATCCATTAAGACCTTTCTTTG | Domestication of *BvDODA1-HA* |
| Silentmut_BvDODA1_R | AGAAAGGTCTTAATGGATGTGTATCTTCTACTGTTAATATTGGGTTTCC | Domestication of *BvDODA1-HA* |
| BbsI_BvDODA1_GG_F | ATGCGAAGACAAGCTTAGTAGTGGAGGTGGATCCATGAAAATGATGAATG | Generation of *BvDODA1-HA* level 0 part |
| BbsI_BvDODA1-HA_GG_R | ATTAGAAGACAATACCTTATGCGTAGTCCGGGACGTCGTACGGATAGGCTGAAGTGAAC | Generation of *BvDODA1-HA* level 0 part |
| BbsI_aroG_F | GCGCGAAGACAAAATGAATTACCAAAATGATGACCTGAGAATTAAAGAAATTAAAGAGTTACTGCCAC | Generation of *aroG*-His* level 0 part |
| BbsI_aroG_R | ATTAGAAGACAAAAGCTTAATGGTGATGGTGATGATGCCCACGACGGGCTTT | |
| BbsI_RhaS_RP | CTCAGAAGACAAAAGCTTATTGCAGAAAGCCATCC | Generation of *rhaS* part for pSH023 |
| BbsI_RhaS_FP | GCGCGAAGACAAAATGACCGTATTACATAGTGTG | |
| BbsI_RhaS_F | ATGCGAAGACAAGCTTAGTAGTGGAGGTTAGCTTATGACCGTATTACA | Generation of *rhaS* part for pSH018 |
| BbsI_RhaS_R | CTCAGAAGACAATACCTTATTGCAGAAAGCCATCCCGTCCCT | |
| Mid_CYP76_SeqR | TCTCGCCCATAGTGAGC | Colony PCR of transformants carrying pSH018 or pSH023 |
| CDS1_SeqF | CGAGTCAGTGAGCGAG | Colony PCR (all plasmids) |
| CDS1_SeqR | CTGACGTCTAAGAAACCATT | Colony PCR of transformants carrying pPMQAK1-T |
